# HIV-1 Vif protein is stabilized by AKT-mediated phosphorylation to enhance APOBEC3G degradation

**DOI:** 10.1101/2021.09.28.462184

**Authors:** Rameez Raja, Chenyao Wang, Akhil C Banerjea

**Author notes:** **Correspondence:** Rameez Raja, or, Akhil C. Banerjea, Mailing address: Department of Inflammation and Immunity (NE4), Cleveland Clinic, 9500 Euclid Avenue, Cleveland, Ohio, 44195, USA.

## Abstract

HIV-1 virus has to counter anti-viral restriction factors for its successful replication after its entry in the cell. The host-pathogen dynamics operate as soon as HIV-1 interacts with the cell. HIV-1 Vif has been known for its role in degradation of APOBEC3G; a cytosine deaminase which leads to hyper mutations in the viral DNA leading to aberrant viral replication. The cellular proteins regulating the intracellular HIV-1 Vif protein levels can have profound impact on HIV-1 pathogenesis. MDM2 is known to induce degradation of Vif with subsequent effects on APOBEC3G. Here, we have identified AKT/PKB as one of the crucial regulators of HIV-1 Vif protein. The rationale for selecting Vif as a target substrate for AKT was the presence of RMRINT motif in it, which is similar to the AKT phosphorylation motif RxRxxS/T. Immunoprecipitation assay and Kinase assay revealed that AKT and Vif interact strongly with each other and Vif is phosphorylated at T20 position by AKT. This phosphorylation stabilizes HIV-1 Vif while Vif mutant T20A degrades faster. Moreover, use of dominant negative form of AKT (KD-AKT) and AKT inhibitors were found to destabilise Vif and increase its K48-ubiquitination profile. The consequences of this AKT-Vif interplay were also validated on APOBEC3G degradation, a target of Vif. AKT inhibition was found to restore APOBEC3G levels. This process can be interpreted as a strategy used by virus to prevent MDM2 mediated Vif degradation; AKT stabilises Mdm2, which then targets Vif for degradation but at the same time AKT stabilises Vif by phosphorylating it. Thus, AKT mediated stabilization of Vif might compensate for its degradation by MDM2. This study can have significant implications as HIV-1 Tat protein and growth factors like insulin activate PI3-K/AKT Kinase pathway and can potentially affect Vif and APOBEC3G protein levels and hence HIV-1 pathogenesis.

## Introduction

HIV-1 is a small retrovirus which manipulates the host proteins to survive and replicate inside the host by exploiting its cellular machinery (1, 2). HIV-1 Vif plays important role in HIV-1 survival in the host cell by counteracting the restriction factor, APOBEC3G. APOBEC3G induces hypermutations in viral DNA by cytidine deaminase activity leading to degradation of viral DNA. If hypermutated viral DNA gets integrated in the cellular genome, it cannot code for functional viral proteins (3). Vif recruits E3 ubiquitin ligase complex and decreases the levels of APOBEC3G protein via proteasomal degradation pathway (4, 5). Vif deficient viruses are severely compromised and unable to multiply in host cells. The cellular proteins regulating the Vif activity can have profound effect on HIV-1 pathogenesis. MDM2, an E3 ligase has been shown to interact with Vif leading to its ubiquitination followed by its proteasomal degradation which results in an increase in levels of APOBEC3G. Core Binding Factor β (CBF β) is known to stabilize Vif and hence counteract anti-viral effect of APOBEC3G (6, 7). On the other hand, apoptosis signal-regulating kinase-]1 (ASK-1) disrupts the interaction between Vif and APOBEC3G to restore the anti-viral activity of APOBEC3G (8).

HIV-1 Tat protein is already known to play important in activation of PI3/AKT signaling pathway (9, 10). Mdm2 is a downstream target of AKT(11). Previously, we have shown that HIV-1 Tat protein stabilizes Mdm2 by inducing its phosphorylation in AKT dependent manner MDM2 is known to enhance the Tat mediated LTR activity (10). MDM2 ubiquitinates Tat at lysine 71 position to potentiate its activity in a non-proteolytic way (12).Thus, there is a positive feed-back loop between Tat, AKT and Mdm2. As, Mdm2 also ubiquitinates HIV-1 Vif protein and induces its proteasomal degradation, so we were interested to study the role of AKT in regulating the HIV-1 Vif levels. The other interesting feature for selecting Vif was the presence of AKT phosphorylation motif RMRINT (RXRXXS/T); similar motifs have been found in AKT target substrates like FKRHL1, IKKα and P21 (11, 13). AKT mediated phosphorylation in these substrates protein regulates their function. We show that AKT stabilizes Vif protein expression and leads to APOBEC3G degradation. This study can have significant implications towards a better understanding of HIV-1 pathogenesis.

## Materials and methods

### Cell culture and transfection

HEK-293T (Human Embryonic Kidney 293T cells) were maintained in Dulbecco’s Modified Eagle’s Medium (DMEM; Himedia Laboratories) supplemented with 10% fetal bovine serum (Gibco, Invitrogen), 100 units penicillin, 0.1mg streptomycin and 0.25μg amphotericin B per ml at 37°C in presence of 5% CO_2_ in a humidified incubator. Transfections were performed using Lipofectamine 2000 (Invitrogen, USA) and Polyethyleneimine, Linear (MW 25,000, Polysciences Inc., USA) reagents using the manufacturer’s protocol.

### Plasmid constructs and chemicals

Viral genes were obtained as previously described (14). HA-APOBEC3G was cloned using cDNA from TZMbl cells in pCMV-HA vector from Clontech, USA. pBlue3’LTR-luc was obtained from NIH AIDS Reference and Reagent Program of NIH, MD, USA. GST Tat was generated by cloning pNL4-3 derived *tat* gene in pGEX-4T1 vector from Addgene. HA Tat and Flag NQO1 were purchased from Addgene. HA Mdm2 was purchased from Sino Biologicals, USA. Myc -Vif (T20A) was made from wild type by site directed mutagenesis. HA-AKT, HA-KD-AKT (K179A), HA-Myr-AKT, GST AKT were a kind gift from Hui Kuan Lin, MD Anderson cancer centre, Texas. Renilla luciferase plasmid was a kind gift from Vivek Natrajan, IGIB, Delhi, India. His Ub plasmid was gifted by Dimitris Xirodimas, University of Dundee. Chemicals used were AKTi (Sigma), IPTG (Sigma) and Insulin (Sigma), Cycloheximide (Sigma) and MG132 (Sigma).

### Western analysis

HEK-293T cells were transfected with gene of interest for 36 hrs. The the cells were harvested and lysed in RIPA lysis buffer (1% NP-40, 20mM TrisCl, pH 7.5, 150 mM NaCl, 1mM Na_2_EDTA, 1mM EGTA, 1% Sodium deoxycholate, 1mM Na_3_VO_4_). Protein estimation was carried out using BCA Protein Assay Kit (Pierce, Thermo Scientific, USA). An equal amount of protein was loaded on SDS-PAGE and was transferred to nitrocellulose membrane. The membranes were blocked with 5% non-fat dry milk (Himedia Laboratories, India). The primary antibodies used were anti-AKT, anti-GAPDH, anti-phospho-AKT (S473) (Cell Signaling Technology), anti-myc, anti-HA (Clontech), anti-GST (Santa Cruz Biotechnology). The secondary antibodies used were anti-rabbit/mouse-HRP conjugated (Jackson Immuno Research). Blots were developed using ECL (Enhanced Chemiluminescence) reagent.

### Cycloheximide chase assay

To study the degradation kinetics of proteins, cycloheximide chase assay was performed. HEK-293T cells were transfected with gene of interest for 24 hrs and treated with CHX (100μg/ml; Sigma). Cell lysates were prepared at indicated time points and subjected to 10% SDS-PAGE followed by western analysis as described above.

### *In vivo* ubiquitination assay

*In vivo* ubiquitination assay was performed to detect ubiquitylated proteins in transfected mammalian cells as described earlier (15). HEK-293T cells were co-transfected with plasmid encoding desired gene and His-Ub (6X Histidine-ubiquitin) for 36 hrs. After 36 hrs of transfection, 25μM of MG132 (Sigma-Aldrich) was added and the cells were further incubated for 8 hrs. The cells were lysed in Buffer A (6M guanidinium-HCl, 0.1M Na_2_HPO_4_/NaH_2_PO_4_, 10mM imidazole; pH 8.0), sonicated, and centrifuged. Ni-NTA beads were added to the supernatant and the mixture was incubated at room temperature for 6 hrs while rotating. Subsequently, the beads were washed with buffer A and buffer TI (25mM Tris, pH 6.8, 20mM imidazole). The ubiquitinated proteins were eluted in buffer containing 200mM imidazole, 5%SDS, 0.15M Tris, pH 6.7, 30% glycerol, and 0.72M β-mercaptoethanol. The eluates were resolved by SDS-PAGE followed by western analysis.

### Site directed mutagenesis and sequencing

To generate Myc Vif with mutation in RMRINT motif, site directed mutagenesis was carried out by using Quik Change II Site-Directed Mutagenesis Kit (Agilent, USA). The primers used were: 5’ GTAGACAGGATGAGGATTAACGCCTGGAAAAGATTAGTAAAACAC 3’ forward primer) & 5’ GTGTTTTACTAATCTTTTCCAGGCGTTAATCCTCATCCTGTCTAC 3’ (reverse primer). The PCR conditions followed were: 95°C (1 minute), 95°C (50 seconds), 60°C (50 seconds), 95°C (5 minutes), 95°C (7 minutes). After thermal cycling, Dpn I treatment was given to digest parental and hemi-methylated DNA followed by the transformation in competent cells. The sequencing was done by SciGenome Labs, India. The clone positive for the mutation as shown by sequencing results was used for the experiments.

### GST-AKT expression and purification

pGEX-4T1-AKT was transformed in BL21 strain of *E*.*Coli* for expression and subsequent purification of GST-AKT. The bacterial culture was induced with 0.5 mM IPTG for 16 hours at 16°C. The cells were lysed by adding lysozyme (1mg/ml) at 4°C with gentle shaking. DTT was added to the bacterial lysate after lysozyme treatment (100μl of 1M DTT). This was followed by sonication and extraction of proteins with Triton X-100. The solution was centrifuged at 12,000 rpm for 15 minutes at 4°C. The supernatant was then used for binding to glutathione beads at 4°C for 3 hours. The beads were centrifuged at 2500 rpm for 2 minutes at 4°C.The beads were washed till the supernatant stops giving colour using Bradford reagent. A control with GST only, bound to glutathione beads was expressed and purified in similar way.

### GST-Pull down assay

GST alone and GST-AKT bound to glutathione beads were expressed and purified as described above. HEK-293T cells were transfected with 2μg of Myc-Vif for 36 hours. The cells were lysed in RIPA buffer. 10ug of GST-AKT was incubated with the cell lysate for 4 hours at 4°C. After incubation, the supernatant was discarded and the beads were washed 5-6 times with chilled 1×PBS. The beads were boiled in laemmli buffer and subjected to SDS-PAGE followed by immunoblotting with anti-Myc and anti-GST antibodies.

### *In vitro* AKT kinase assay

HA AKT and its substrate proteins were transfected in HEK-293T cells. These proteins were purified by immunoprecipitation. Then purified AKT (0.5μg) was added to the purified substrate protein (2μg) along with AKT kinase reaction buffer (50mM HEPES, 0.01% Tween 20, 10mM MnCl2, 1mM EGTA, 2.5mM DTT and 0.1mM ATP, pH 7.4) for 60 minutes at 30°C. All reactions were stopped by adding 5X SDS loading buffer and boiled for 10 min at 95°C for western blotting analysis.

### Statistical analysis

Results obtained were represented as mean±standard error of the mean (s.e.m). *P-values* were calculated by a two-tailed t-test. Only values with p<0.05 were considered significant.

## Acknowledgements

Several reagents were obtained from AIDS Reference & Reagent Program of NIH, MD, USA. This work was supported by Department of Biotechnology and Department of Science and Technology of Government of India.

## Author contributions

Rameez Raja and Akhil C Banerjea conceived the idea and designed experiments. Rameez Raja and Chenyao Wang performed the experiments. Rameez Raja and Akhil C Banerjea wrote the manuscript.

## Results

### Screening of viral genes as a target substrate for AKT

To analyze the effect of AKT on expression of HIV-1 Vif and other viral accessory genes, Myc-tagged viral genes were transfected in HEK-293T cells. After 36 hours of transfection, AKT inhibitor (AKTi) was added for 8 hours and then the cells were harvested. The cells were lysed and analyzed by immunoblotting for the expression levels of different viral proteins (Fig. 1a, 1b, 1c, 1d and 1f). From our screening experiments, we identified that the expression of two viral proteins namely, HIV-1 Tat and HIV-1 Vif was reduced in presence of AKT inhibitor (Fig. 1c and 1f). These data suggest that AKT regulates Tat and Vif which can have significant consequences on of Tat-AKT-Mdm2 axis and hence HIV-1 pathogenesis.

**Figure 1.**
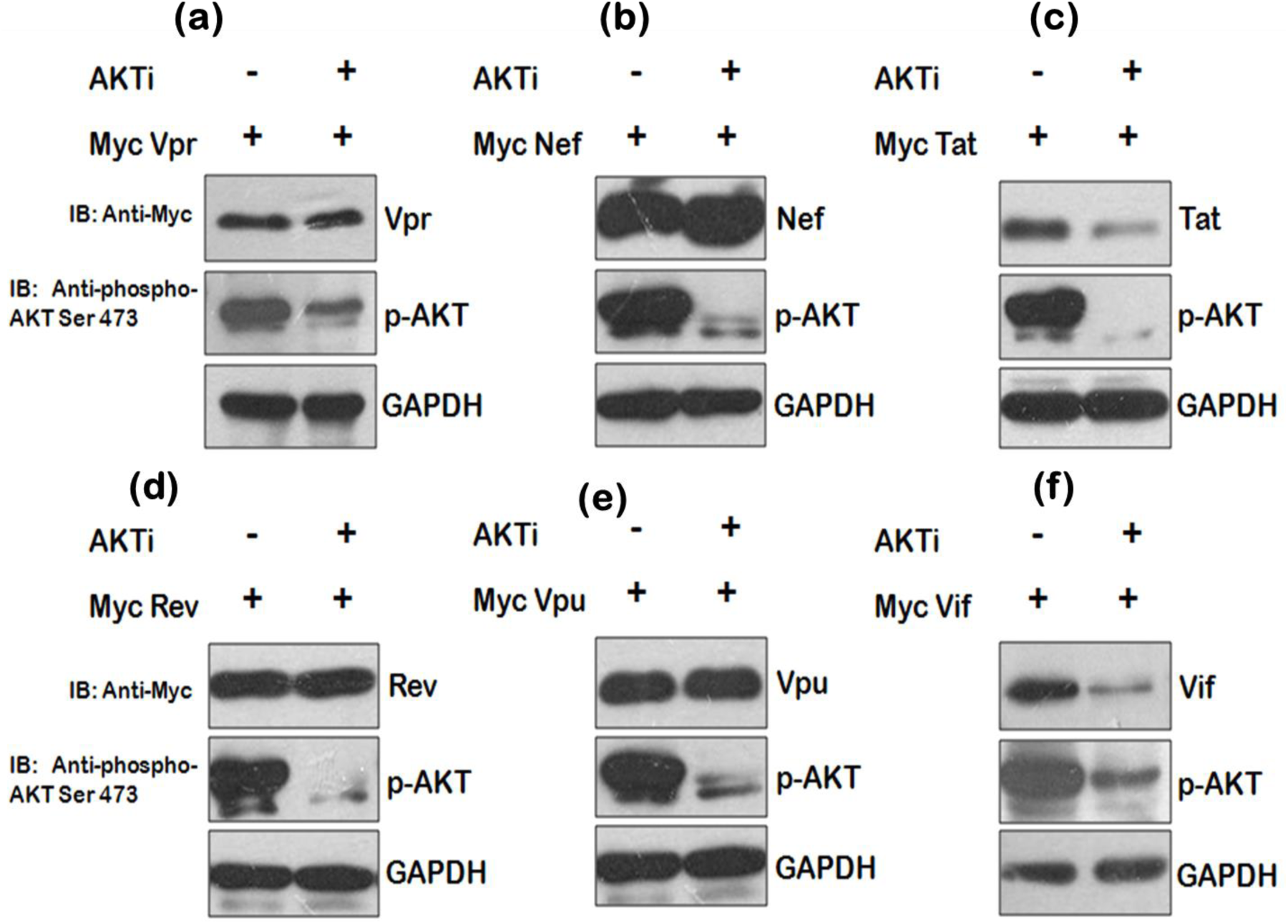
Screening of HIV-1 genes. **(a)** HEK-293T cells were transfected with Myc-Vpr, Myc-Nef, Myc-Tat, Myc-Rev, Myc-Vpu and Myc Vif plasmids for 36 hrs. AKT inhibitor treatment was given for 8 hrs before harvesting cells. Cell lysates were analyzed by western blotting using anti-Myc, anti-p-AKT and anti-GAPDH antibodies. GAPDH was used as loading control.

### AKT inhibition decreases HIV-1 Vif protein levels in a dose dependent manner

As HIV-1 Vif was having an AKT phosphorylation motif RMRINT, here we studied its regulation in more detail (Fig. 2a). However, the consequences of AKT mediated Tat regulation remain to be unexplored. The effect of AKT on Vif levels was also investigated by co-transfection of HA-Myr-AKT (constitutively active form of AKT) along with Myc-Vif in HEK-293T and MCF-7 cells. The cells were harvested after 36 hours of transfection followed by immunoblotting. A significant increase in Vif levels was observed in presence of Myr-AKT in both these cell lines (Fig.2b, 2c). The expression of Vif was significantly reduced in the presence of HA-KD-AKT indicating that AKT activity controls Vif expression (Fig. 2d). HIV-1 Vif protein expression was also investigated in presence of different concentrations of AKTi at 2μM and 4μM. Both the doses reduced the expression of HIV-VIf protein with 4μM dose showing maximal effect (Fig. 2e, 2f). The effect of dose dependent inhibition of AKT activity was observed more prominently by using KD-AKT (kinase deficient AKT), a dominant negative form of AKT. Myc-Vif was co-transfected along with different concentrations of HA-KD-AKT in HEK-293T cells. KD-AKT was able to decrease the expression level of HIV-1 Vif protein in a dose dependent manner (Fig.2g). Thus, these data indicate that AKT regulates the levels of Vif protein.

**Figure 2.**
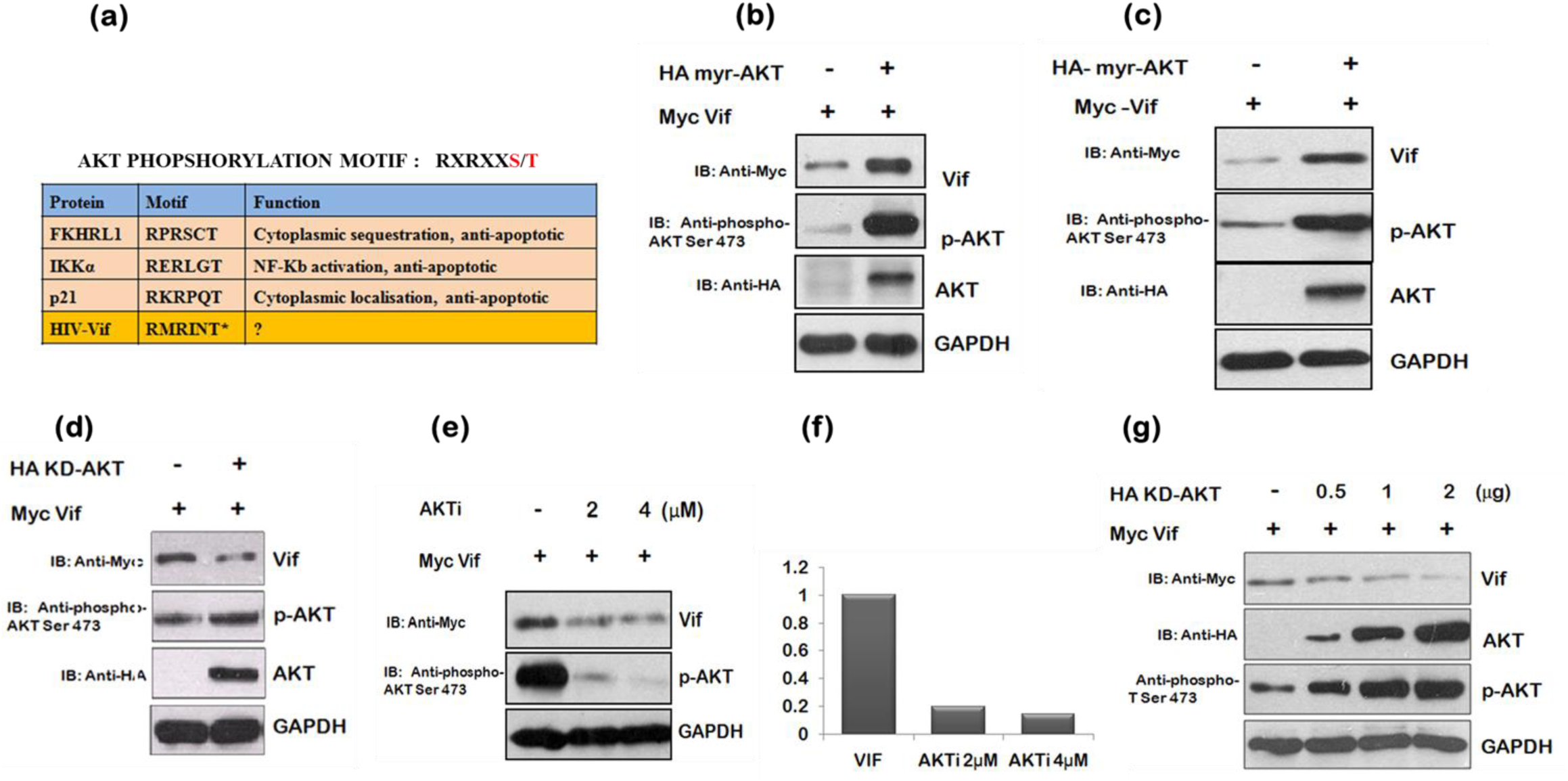
AKT inhibition decreases HIV-1 Vif protein levels in a dose dependent manner. **(a**) Table showing AKT phosphorylation motif in different proteins and HIV-1 Vif protein. **(b)** HEK-293T cells were co-transfected with Myc-Vif and HA-KD-AKT as indicated. After 24hrs of transfection, cells were lysed and cell lysates were subjected to western blotting using anti-HA, anti-phospho-AKT Ser473, anti-Myc and anti-GAPDH antibodies. GAPDH was used as loading control. **(c)** MCF-7 cells were co-transfected with Myc-Vif and HA KD-AKT as indicated. After 24hrs of transfection, cells were lysed and cell lysates were subjected to western blotting using anti-HA, anti-phospho-AKT Ser473, anti-Myc and anti-GAPDH antibodies. GAPDH was used as loading control. **(d)** HEK-293T cells were transfected with Myc-Vif for 24hrs followed by AKTi treatment of 2 μM and 4 μM for 8 hours. Cells were lysed and cell lysates were subjected to western blotting using anti-HA, anti-phospho-AKT Ser473, anti-Myc and anti-GAPDH antibodies. GAPDH was used as loading control. **(e)** Graph showing the densitometry analysis for Fig. 1d. **(f)** HEK-293T cells were transfected with Myc-Vif along with different concentrations of HA-KD-AKT for 24hrs. Cells were lysed and cell lysates were subjected to western blotting using anti-HA, anti-phospho-AKT Ser473, anti-HA, anti-Myc and anti-GAPDH antibodies. GAPDH was used as loading control.

### AKT regulates Vif expression at post-translational level

After we found that HIV-1 Vif protein is a target for AKT, we inquired whether the effect is at transcriptional or post translational level. For this, we performed cycloheximide chase assay to find out the mechanism of AKT mediated vif regulation. Cycloheximide inhibits new protein synthesis and hence is used to assess the stability of protein at post-translational level. Myc-Vif was transfected either alone or along with with HA-Myr-AKT in HEK-293T cells. In presence of Myr-AKT, Vif expression levels were observed to be more stable even after longer time duration of CHX treatment (Fig.3a). These results suggest that Vif expression is regulated by AKT at post-translational level.

**Figure 3.**
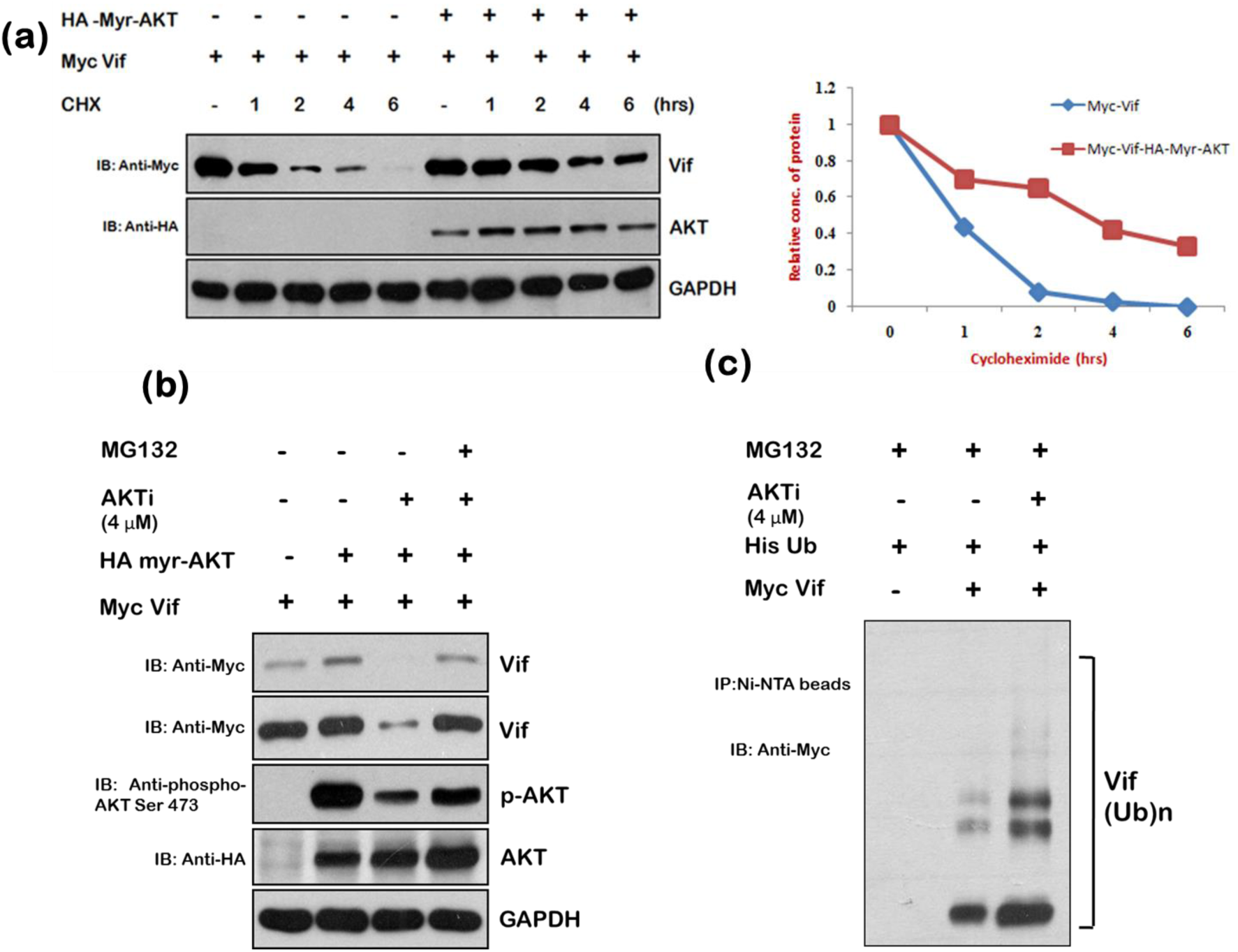
AKT stabilises Vif expression at post-translational level. **(a)** HEK-293 T cells were transfected with Myc-Vif either alone or along with HA-Myr-AKT for 36 hrs. Cells were treated with Cycloheximide (100μg/ml) for the indicated time periods. Cell lysates were subjected to western blot analysis using anti-HA, anti-Myc and anti-GAPDH antibodies. GAPDH was used as loading control. Densitometric analysis was done by ImageJ and shown as line graph. **(b)** HEK-293T cells were co-transfected with HA-Myr-AKT and Myc-Vif plasmids and treated with AKTi (4μM) for 24 hrs. After 24 hrs, cells were treated with MG132 (10μM) for 8hrs. Cell lysates were analyzed by western blotting using anti-HA, anti-Myc, anti-phospho-AKT Ser473 and anti-GAPDH antibodies. GAPDH was used as loading control. **(c)** HEK-293T cells were transfected with Myc Vif and His Ub expression plasmids and treated with AKTi (5μM) for 24 hrs. Cells were treated with MG132 (10μM) for 8hrs. Cell lysates were subjected to immunoprecipitation with Ni-NTA beads followed by western blotting using anti-Myc antibody.

In order to assess the involvement of proteasomal pathway in Vif protein regulation by AKT, HEK-293T cells were transfected with Myc-Vif alongwith Myr-AKT. After 36 hours of transfection cells were treated for 8 hours with AKTi either alone or with MG-132 (proteasomal inhibitor). The cells were harvested and subjected to immunoblotting to analyze the Vif protein expression levels. As expected, Vif levels were increased in presence of Myr-AKT and decreased in AKTi treated cells. Interestingly, the Vif degradation was rescued in presence of MG132 treated cells, indicating the role of proteasomal degradation pathway in the process (Fig.3b).

As proteasomal pathway was found to play a role in AKT mediated regulation of Vif protein levels, we also assessed the ubiquitination levels of Vif protein in presence of AKTi with MG132 treatment in all samples. MG132 blocks proteasome pathway and prevents K48 ubiquitinated proteins from degradation. HEK-293T cells were transfected with Myc-Vif along with His–Ub with or without AKTi treatment. The cells were treated with MG132 for 8 hours and the ubiquitinated species were detected by immunoblotting after pull down with Ni-NTA beads. The ubiquitinylated Vif increased in presence of AKTi indicating post-translational regulation of Vif by AKT (Fig.3c).

### HIV-1 Vif interacts with AKT and is phosphorylated by it

To analyze the interaction of HIV-1 Vif with AKT, GST-AKT was purified from the bacterial expression system using BL21 cells as shown in Fig. 4a. 10μg of GST-AKT was used for binding to gluthaione beads and subjected to immunoprecipitation by using Myc-Vif-transfected cell lysates and GST alone or GST-AKT conjugated to glutathione beads. GST-AKT showed a strong interaction with Vif protein as shown in the (Fig.4b). AKT, being a kinase, is known to phosphorylate its target proteins having consensus AKT phosphorylation motif RXRXXS/T. HIV-1 Vif has AKT phosphorylation motif RMRINT in its sequence which is similar to the consensus motif and interacted with AKT. Next we performed Kinase assay to validate the role of this phosphorylation motif. The Threonine residue in the HIV-1 Vif phosphorylation motif RMRINT was mutated to T20A from Myc-Vif wild type by site directed mutagenesis (Fig. 4c) and was used a control to validate the role of threonine residue it phosphorylation. Indeed, we found that HIV-1 Vif Wt was phosphorylated by AKT while the HIV-1 Vif T20A was not (Fig.4d). These data indicate that HIV-1 Vif is phosphorylated by AKT at T20 position.

**Figure 4.**
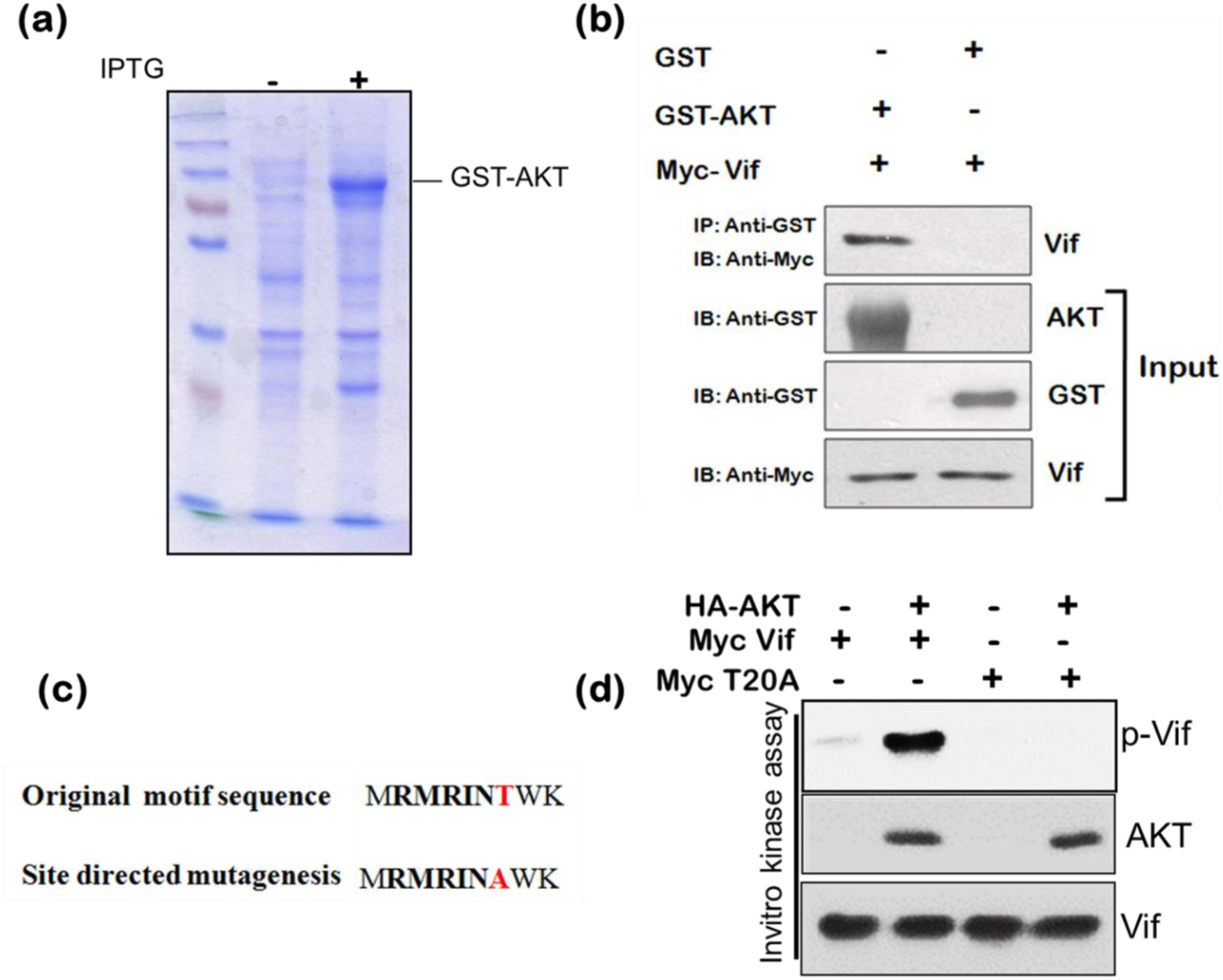
HIV-1 Vif interacts with AKT and is phosphorylated by it. **(a)** GST-AKT was purified from bacterial system using IPTG induction as described in materials and methods. **(b)** HEK-293T cells were transfected with Myc-Vif encoding plasmid. After 24 hrs of transfection, cell lysates were prepared and incubated with GST-AKT fusion protein and GST alone bound to GST beads for 24 hrs at 4°C. The bound proteins were analyzed by western analysis using anti-Myc antibody. GST alone was used as negative control. **(c)** HIV-1 Vif has RMRINT motif (AKT phosphorylation motif). The RMRINT point mutant Myc-Vif (T20A) was generated using site-directed mutagenesis. **(d)** HEK-293T cells were transfected with HA-AKT, Myc-Vif and mutant Myc-Vif (T20A). AKT and Vif proteins were purified by immunoprecipitation. The purified AKT was added to the purified Vif and its mutant (T20A) along with kinase reaction buffer. The phosphorylated AKT substrates were analyzed by western blotting.

### Stability of Vif mutant T20A is less than wild type

HEK-293 T cells were transfected with either Myc-Vif wild type or mutant Myc-Vif (T20A). The stability of both the proteins was assessed by cycloheximide chase assay. After 36 hours of transfection, CHX was added to the cells. The cells were lysed at different time points and analyzed by immunoblotting using anti-Myc antibody. The mutant Vif (T20A) showed faster degradation kinetics compared to wild type (Fig. 5a,5b). This result further confirms the role of AKT in mediating Vif stabilization.

**Figure 5.**
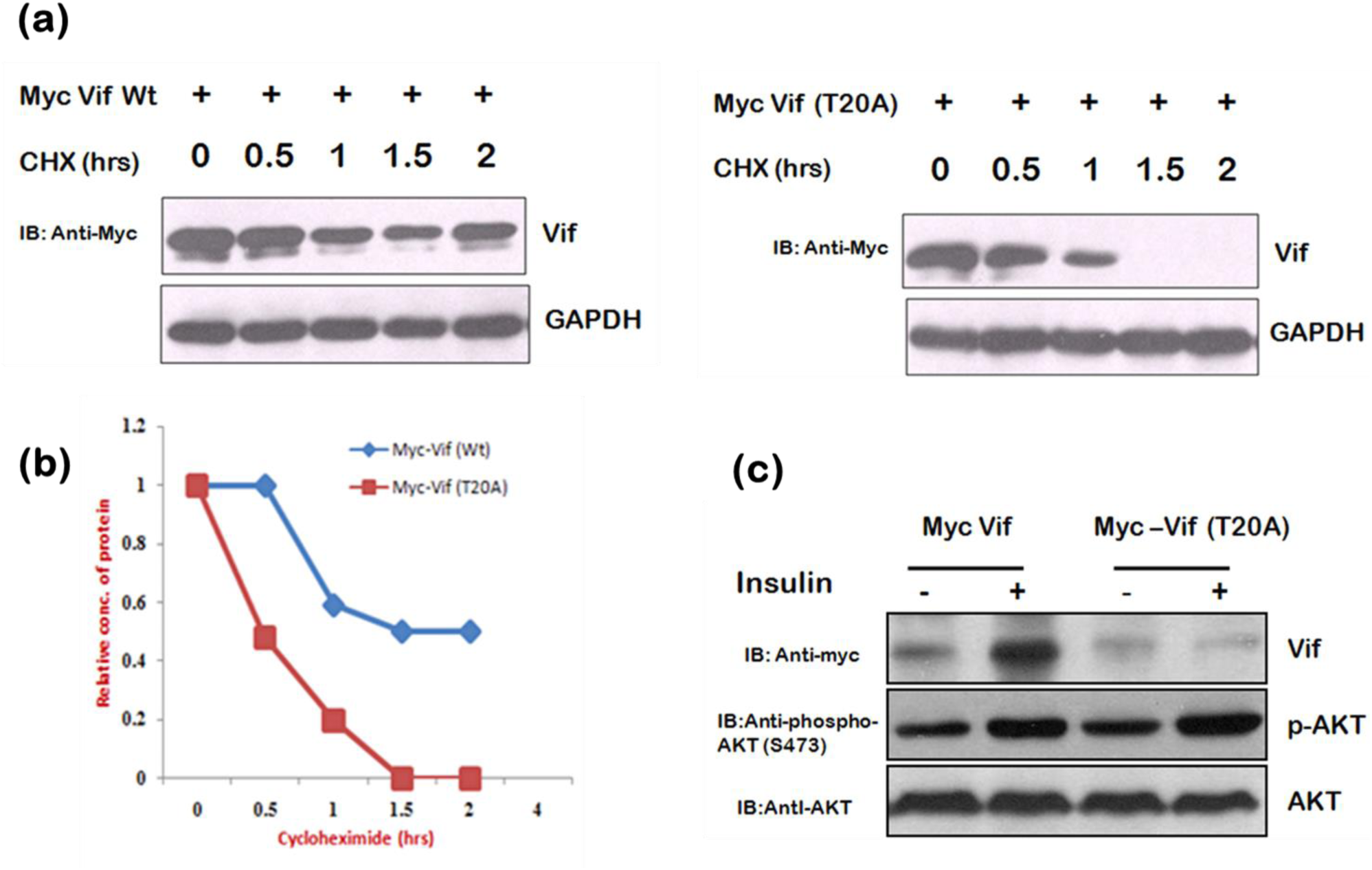
Stability of Vif mutant T20A is less than wild type. **(a, b)** HEK-293T cells were transfected with Myc Vif or Myc Vif (T20A) expression plasmids for 36 hrs and treated with CHX (100μg/ml) for indicated time periods. Cell lysates were analyzed by western blotting with anti-Myc and anti-GAPDH antibodies. GAPDH was used as loading control. Densitometric analysis was done by ImageJ and shown as line graph control. **(c)** HEK-293T cells were transfected with Myc-Vif and Myc-Vif (T20A) expression plasmids as indicated for 36 hrs. Cells were treated with Insulin and cell lysates were subjected to western blotting analysis using anti-Myc, anti-phospho-AKT Ser473 and anti-AKT antibodies. AKT was used as internal loading.

### Insulin increases levels of Vif but not Vif (T20A) mutant

Insulin is a growth factor which stimulates PI3/AKT signaling pathway. It was used to validate the above finding. HEK-293T cells were transfected with Myc-Vif and Myc-Vif (T20A) mutant. After 36 hrs of transfection, cells were treated with insulin. The cells were lysed and subjected to immunoblotting. HIV-1 vif expression was found to be increased in the presence of insulin while the levels of Vif (T20A) mutant were not affected by insulin treatment (Fig. 5c). This confirmed our finding that AKT positively regulates the expression levels of HIV-1 Vif.

### Down-regulation of Vif by AKT inhibition restores APOBEC3G levels

The functional consequences of the AKT mediated stabilization of Vif were investigated by using the known target of Vif. The Vif protein is known to induce proteasomal degradation of APOBEC3G, an anti-viral restriction factor. HEK-293T cells were transfected with HA-APOBEC3G alone or along with Myc-Vif. The cells were treated with 4μM AKTi after 36 hours of transfection. The cells were lysed and analyzed by immunoblotting to assess the levels of APOBEC3G. The levels of APOBEC3G were reduced significantly in presence of Myc-Vif compared to APOBEC3G alone. However, the cells treated with AKTi showed reversion in APOBEC3G levels even in presence of Vif. The reversion was seen due to AKTi mediated down regulation of Vif (Fig.6a). The results were validated with KD-AKT also. HEK-293 T cells were transfected with HA-APOBEC3G alone or along with Myc-Vif and HA-KD-AKT. The cells were collected after 36 hours of transfection and subjected to immunoblotting. KD-AKT was found to be able to restore the APOBEC3G levels in presence of Vif as it decreases Vif protein levels (Fig.6b).

**Figure 6.**
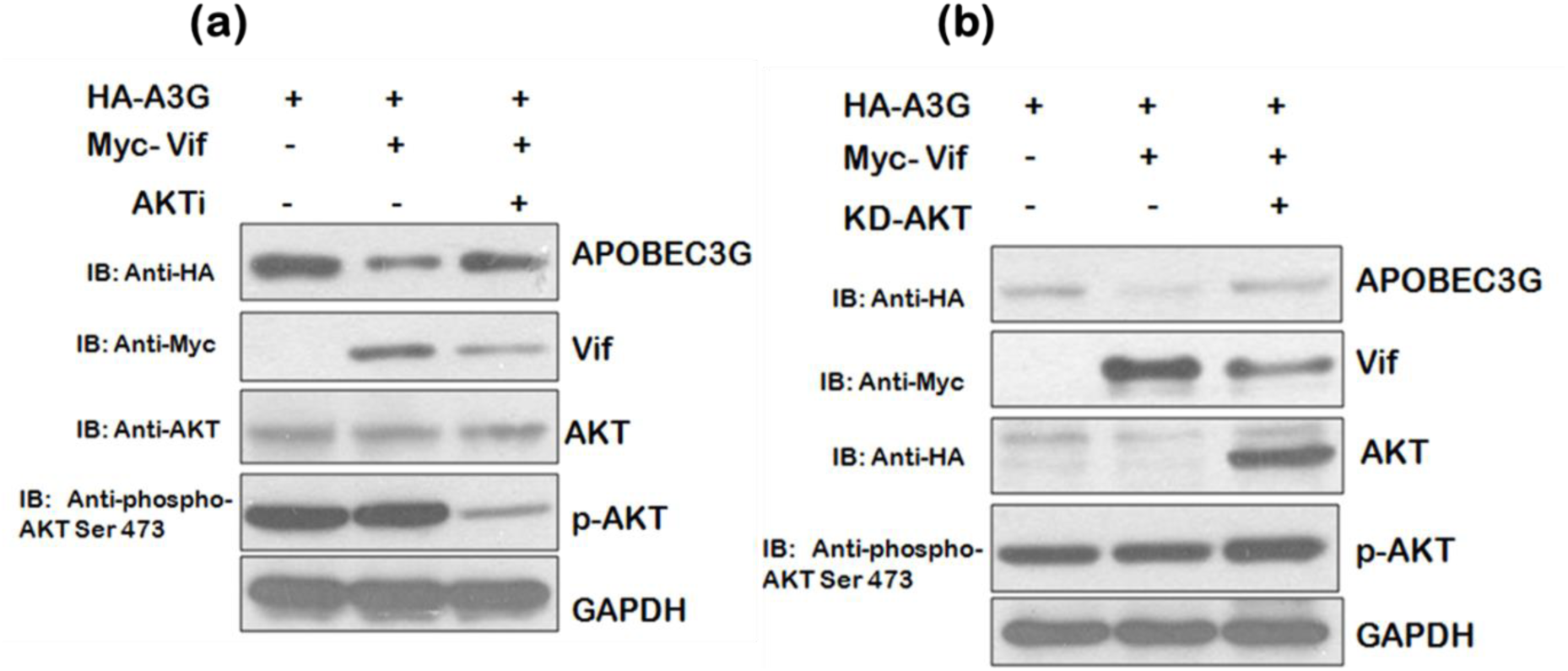
Down-regulation of Vif by AKT inhibition restores APOBEC3G levels. **(a)** HEK-293T cells were co-transfected with HA-APOBEC3G and Myc-Vif as indicated and treated with AKTi (4μM) after 36 hrs of transfection. Cell lysates were subjected to SDS-PAGE followed by western blotting using anti-HA, anti-myc, anti-phospho-AKT Ser473, anti-AKT and anti-GAPDH antibodies. GAPDH was used as loading control. **(b)** HEK-293T cells were co-transfected with HA-APOBEC3G, Myc Vif and HA KD-AKT as indicated for 36 hrs. Cells were lysed and cell lysates were analyzed by SDS PAGE followed by western blotting using anti-HA, anti-myc, anti-phospho-AKT Ser473 and anti-GAPDH antibodies. GAPDH was used as loading control.

## Discussion

There are number of restriction factors which protect the host cells from the pathogen. HIV-1 has evolved various mechanisms to counter these host cell factors. The well-known anti-HIV-1 restriction factors are TRIM5 (tripartite motif 5) (16), APOBEC3G (apolipoprotein B mRNA-editing enzyme-catalytic polypeptide-like 3G) (17, 18), SAMHD1 (19, 20) and Tetherin (or BST-2)(21). TRIM5 inhibits the uncoating of capsid proteins which is required to release viral mRNA into host cell. SAMHD1 depletes the cellular pool of dNTPs to reduce HIV-1 replication. APOBEC3G is cytidine deaminase that converts cytosine to uracil. These mutations lead to the degradation of proviral cDNA and even if the cDNA is integrated into host genome, it is rendered non-functional due to the mutations. Tetherin binds to the surface of cell at the site of budding virions and blocks the release of viral progeny. HIV-1 has developed various strategies to evade these restriction factors except TRIM5 as none of the HIV-1 proteins is known to target this. The restriction factors SAMHD1, Tetherin and APOBEC3G are targeted by Vpx/Vpr, Vpu (or Env) and Vif respectively (3).

HIV-1 Vif recruits E3 ligase complex consisting CUL5, ELOB/C, RBX and CBFβ for APOBEC3G degradation (6, 22). However, there are very few reports about the regulation of Vif protein (7). Core Binding Factor β (CBF β) has been shown to stabilize Vif with subsequent effects on APOBEC3G levels (23). On the other hand, MDM2 is reported to promote Vif degradation to elevate APOBEC3G levels (24). As, Mdm2 is a downstream target of AKT and inter-regulation of HIV-1 Tat, AKT and Mdm2 is already known, we investigated the role of AKT in regulation of HIV-1 Vif. Inhibition of AKT activity by using AKTi or dominant negative form of AKT i.e. KD-AKT was found to reduce the expression level of Vif. The constitutively active form of AKT i.e Myr-AKT significantly increased the expression of Vif protein. Also, AKT and Vif were found to interact strongly with each other in GST pull down assay. The stability of Vif in presence of Myr-AKT was also observed to be more in cycloheximide chase assay. To investigate the mechanism of this regulation, ubiquitination assay was carried out which showed an increase in the levels of ubiquitinylated Vif in presence of AKTi. MG132 treatment was done to confirm the involvement of proteasomal pathway in the Vif degradation. It also showed that MG132 treatment reverses the effect of AKTi on the levels of Vif. Further, the functional validation of AKT mediated Vif stabilization was carried out by analyzing APOBEC3G expression levels, a downstream target of Vif, in a co-transfection system in presence of AKTi or KD-AKT. The APOBEC3G levels were also reverted by downregulation of Vif either by AKTi or KD-AKT. The mutation of Threonine to Alanine in the AKT phosphorylation motif RMRINT decreased the stability of Vif significantly as observed in cycloheximide chase assay. We also observed that insulin treatment leads to the enhancement in the levels of Vif protein as it is a growth factor and activates AKT signaling pathway resulting in the phosphorylation and activation of AKT. These results suggested that AKT increases the levels of HIV-1 Vif protein and helps the virus in combating APOBEC3G

The exploitation and modulation of host cellular signaling pathways by HIV-1 proteins is a very complex phenomenon. HIV-1 Tat is an early protein of the virus which activates AKT signaling pathway (9, 10). On the one hand, AKT phosphorylates MDM2 resulting in its stabilization as previously shown by us and MDM2 further increases the LTR transcriptional activity of Tat by inducing its non-proteolytic K63 ubiquitination thus creating a positive feedback loop between Tat, AKT and MDM2 (10, 12). On the other hand, AKT also increases the levels of HIV-1 Vif protein. Vif is ubiquitinated and degraded by MDM2 which is activated by AKT itself. Thus, HIV-1 exploits AKT signaling pathway through its activation by Tat resulting in the enhancement of Vif levels in order to overcome the degradation of Vif by MDM2 (Fig.7). It helps the virus to evade the host restriction factor APOBEC3G more effectively and survive in the host cell.

**Figure 7.**
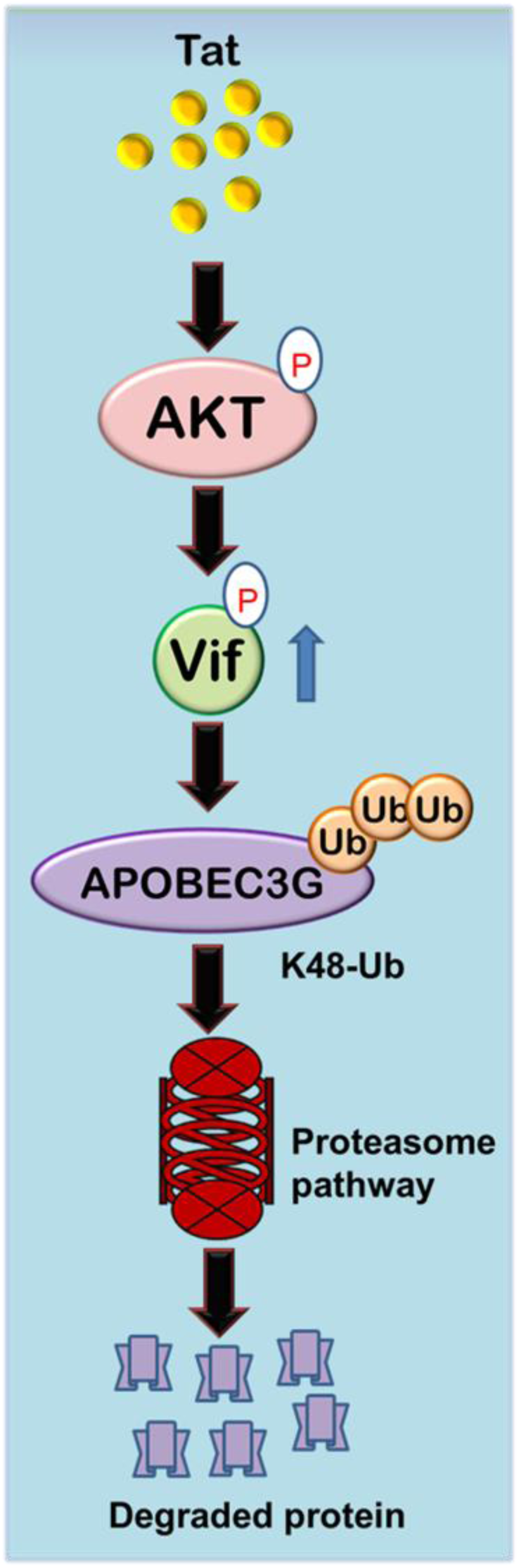
Mechanistic Model showing Tat-AKT-Vif axis to regulate APOBEC3G levels. HIV-1 Tat activates AKT and activated AKT in turn phosphorylates Vif to stabilize it, which then targets APOBEC3G for degradation to promote viral replication.

